# Continuous flash suppression and monocular pattern masking impact subjective awareness similarly

**DOI:** 10.1101/264598

**Authors:** J.D. Knotts, Hakwan Lau, Megan A. K. Peters

## Abstract

Peters & Lau (2015) found that when criterion bias is controlled for, there is no evidence for unconscious visual perception in normal observers, in the sense that they cannot directly discriminate a target above chance without knowing it. One criticism of that study is that the visual suppression method used, forward and backward masking (FBM), may be too blunt in the way it interferes with visual processing to allow for unconscious forced-choice discrimination. To investigate this question we compared FBM directly to continuous flash suppression (CFS) in a two-interval forced choice task. Although CFS is popular, and may be thought of as a more powerful visual suppression technique, we found no difference in the degree of perceptual impairment between the two suppression types. To the extent that CFS impairs perception, both objective discrimination and subjective awareness are impaired to similar degrees under FBM. This pattern was consistently observed across 3 experiments in which various experimental parameters were varied. These findings provide evidence for an ongoing debate about unconscious perception: normal observers cannot perform forced-choice discrimination tasks unconsciously.

## Introduction

Whether normal observers can perform forced-choice discrimination tasks unconsciously, i.e., whether thresholds for objective performance and subjective awareness on these tasks in normal observers can dissociate, is controversial (Phillips & Block, 2016; Peters et al., 2017; Phillips, 2017). This is an important issue to resolve, because a reliable means of demonstrating unconscious perception would be an invaluable tool for studying the neural correlates of consciousness while controlling for unconscious signal processing confounds (Lau, 2008; Lau & Rosenthal, 2011).

While the dissociation of objective and subjective thresholds is typically thought to occur in blindsight (Weiskrantz, 1986; but see Phillips, 2017 for an opposing view), evidence for the same dissociation in normal observers has been contentious. Some studies have failed to replicate (Kolb & Braun, 1995; Morgan et al., 1997), while others (e.g., Hesselman, Hebart, & Malach, 2011; Salti et al., 2015) have been potentially subject to the well-known confound of criterion bias (Eriksen, 1960; Hannula et al., 2005; Lloyd et al., 2013; Merikle et al., 2001).

A recent study by Peters & Lau (2015) showed that when criterion bias is controlled for using a two-interval forced choice (2IFC) task, there is no evidence for unconscious forced-choice discrimination in normal observers. However, a concern regarding that study is that the method they used for disrupting target visibility, forward plus backward pattern masking (also referred to as sandwich masking, but hereafter referred to as FBM) may interfere at too early a stage in visual processing to facilitate unconscious perception. It could be argued then that if Peters & Lau (2015) had used a visual suppression method that interferes at a later stage, such as metacontrast masking or continuous flash suppression (CFS) (Breitmeyer, 2015), then unconscious perception would have been observed.

We addressed this concern in the current study by directly comparing different visual suppression methods in an adapted version of the 2IFC paradigm used in Peters & Lau (2015). On each trial, a left-or right-tilted target grating in one interval was masked by a monocular pattern masking method (FBM in Experiments 1 & 2, BM in Experiment 3), while a left-or right-tilted target grating in the other interval was masked by a binocular rivalry-based method (CFS in Experiments 1 & 3, interocular suppression (IS) in Experiment 2). Using this setup, if one suppression method is in fact more *permissive* of unconscious processing than the other, then when subjective awareness of the target grating is matched between suppression methods, there should be higher objective discrimination performance under the more permissive method. Similarly, when left-right discrimination performance under the two methods is matched near perceptual threshold, subjective awareness of the target grating should be relatively reduced under the more permissive method. In other words, we should find a difference in the magnitude of any dissociation between objective and subjective discrimination thresholds, or *relative* blindsight (Lau & Passingham, 2006), between the two suppression methods. We tested this hypothesis using a different pair of suppression methods in each of three psychophysical experiments. To anticipate, we did not find such evidence.

## Experiment 1

### Methods

#### Participants

Twenty-five participants (7 female, ages 19-39, 1 left-handed, 9 left-eye dominant), including the first author, gave written informed consent to participate. All participants had normal or corrected-to-normal eyesight and normal stereo vision, and all were either paid $10 USD or given course credit for their participation. The data of five participants were removed due to failure to pass the adaptive staircasing stage (see Procedure section below). The data of one additional participant were removed after they disclosed that they began pushing buttons at random during the main experiment. Therefore, nineteen total participants (6 female, ages 20-39, 1 left-handed, 7 left-eye dominant) were included in the analyses for Experiment 1. This and all subsequent experiments were conducted in accordance with the Declaration of Helsinki and were approved by the UCLA Institutional Review Board.

#### Apparatus and Stimuli

All stimuli were generated with custom Matlab R2013a (Natuck, MA) scripts using PsychToolbox 3.0.12 on a gamma-corrected Dell E773c CRT monitor with a resolution of 1024 x 768 pixels and a 75Hz refresh rate. To achieve binocular rivalry, all stimuli were viewed through a ScreenScope Desktop stereoscope. Target stimuli were sinusoidal gratings with a spatial frequency of .025 cycles/pixel tilted 45° to either the left or the right of vertical. Gratings were 153 pixels in diameter and were viewed through a circular annulus of the same diameter with a Gaussian hull spatial constant of 100. The viewing distance was 33 cm, making grating stimuli approximately 6.5 visual degrees in diameter. Mask stimuli were colored Mondrian patterns of the same dimensions as target stimuli, and were created in Matlab as previously described (Stein, Hebart, & Sterzer, 2011). Target and mask stimuli were presented centered within two square-shaped boxes or “fusion contours” (one for each eye, diameter 7.4°), each side of which was composed of eleven 17×17 pixel squares, alternating between black and white (see Figure 1A). By default, fusion contours were horizontally centered within each left-right half of the screen (11.2 degrees from the midline each) and vertically centered on the screen. At the beginning of each session participants were allowed to shift the on-screen location of the left fusion contours by button press (one pixel per press in any of the cardinal directions), so as to achieve optimal fusion when viewing the screen through the stereoscope. Seven of nineteen participants included in the main analyses used this function (mean ± SD shifts = 3.69° ± 1.97° leftward and 0.31° ± 0.53° downward).

**Figure 1.**
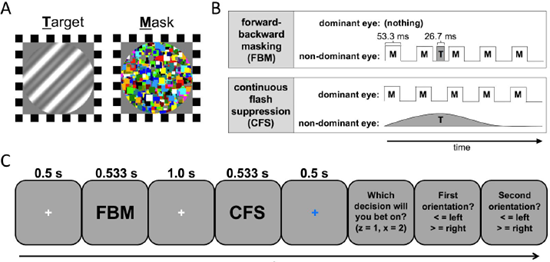
Stimuli and task structure. **A**) Examples of target grating and mask stimuli used in all experiments. **B)** Temporal dynamics of stimuli in Experiment 1. Masks were presented for 53.3 ms each, with intervening blank gaps of the same length. In FBM intervals, masks and target were presented to the non-dominant eye, with the target appearing for 26.7 ms evenly between masks 2 and 3. In CFS intervals, masks were presented to the dominant eye, while the target grating was presented to the non-dominant eye. The contrast of the target ramped up linearly from zero to peak contrast over a period of 173 ms, remained at peak contrast for 26.7 ms, then ramped down linearly to zero over another 173 ms. **C)** Task structure. FBM and CFS stimuli were presented in pseudo random order, separated by a 1.0 s interstimulus interval. Following presentation of stimuli, participants were instructed to bet on the interval in which they felt more confident in their ability to judge the orientation of the target grating. They were then asked to judge the orientations (left or right) of the target gratings in intervals 1 and 2, in that order.

Each trial of the main experiment contained two stimulus intervals: one in which the target was masked by FBM, and the other in which the target was masked by CFS (Figure 1B,C). Each stimulus interval had a total length of 533.3 ms. In both FBM and CFS intervals, a series of 5 different masks was presented to one eye. In the FBM interval, all stimuli were presented to the non-dominant eye. Each mask was presented for 53.3 ms and separated from the next mask by a 53.3 ms blank interval, with the exception of the interval between the 2^nd^ and 3^rd^ masks, in the middle of which the target appeared for 26.7 ms (Figure 1B). The dominant eye was presented with nothing during the FBM interval. In the CFS interval, masks were presented to the dominant eye with the same temporal profile as in the FBM interval. To the non-dominant eye, the target was presented at a range of contrast levels, which started at 0, and ramped up linearly to a peak contrast level over the course of 173 ms. The target stayed at peak contrast for 26.7 ms, and then ramped back down to zero linearly over the course of 173 ms. The last 159.6 ms of the stimulus interval for the non-dominant eye were blank (Figure 1B). Target offset occurred prior to mask offset to prevent image aftereffects that were identified in pilot experiments and have also been identified previously (Tsuchiya & Koch, 2005). The side of target presentation was thus fixed across all trials for each participant to the side of the non-dominant eye. Timing of all stimulus presentations was validated using a Tektronix TDS 3014B oscilloscope.

#### Procedure

The trial structure in the main experiment extends the two-by-two forced-choice (2×2FC) paradigm first introduced by Nachmias and Weber (1975). This method was subsequently used to explore the relationship between detection and identification (e.g., Thomas, Gille, & Barker, 1982; Watson & Robson, 1981), and has more recently been applied to research on perceptual confidence (Barthelemé & Mamassian, 2009, 2010; de Gardelle & Mamassian, 2014). The participant’s task was to discriminate the orientation of a masked target grating (left or right) in each interval (Type 1 decision) and to indicate the interval in which they felt more confident about their orientation judgment (Type 2 decision; Figure 1C). Each trial started with the presentation of a white fixation cross (0.34° diameter) for 0.5 s. This was followed by the two stimulus intervals (described above − 0.5 s each) separated by a 1.0 s inter-stimulus interval containing another white fixation cross. The second stimulus interval was followed by a 0.5 s blue fixation cross to signal the upcoming response period. Participants were then presented with three response prompts, always in the same order, all of which were responded to by button press on a regular computer keyboard. First, participants were asked to make the Type 2 judgment by choosing the interval in which they felt more confident in their orientation judgment. Then participants were asked to make the Type 1 orientation judgments for the targets in the first and second intervals, respectively (Figure 1C). The confidence judgment was placed before the orientation judgments to prevent participants from factoring their reaction times on the orientation task into their confidence judgments. There was no time limit for response, and speed was never emphasized to participants. Participants were also informed that there would be several intervals in which they may not subjectively feel they saw the target, and that for these intervals, they should give their best guess as to the target’s orientation.

Prior to the main experiment, participants completed 42 practice trials. Practice trial structure was identical to that in the main experiment, except for the addition of trial-by-trial feedback about the accuracy of both orientation and confidence responses. A confidence response was considered accurate if the participant bet on a correct orientation judgment. In the first 12 trials, target Michelson contrast was 100% under both suppression conditions. In the first 6 trials, stimuli were displayed at half speed. For the last 30 trials, target contrast was varied independently under each suppression condition according to an adaptive staircase procedure (QUEST, Watson & Pelli, 1983) set to estimate the target stimulus contrast at which orientation discrimination accuracy would be 75% correct. It should be noted, however, that the function of these 30 trials was only to familiarize participants with the task under gradually more difficult conditions. Threshold contrast values were not estimated from practice session data.

Following the practice trials, participants performed another adaptive staircase procedure (QUEST, Watson & Pelli, 1983) to actually estimate the target contrast values at which orientation discrimination accuracy would be matched at 75% correct for both suppression methods. This procedure consisted of 4 blocks of 40 trials each, where the trial structure was identical to that of the main experiment (Figure 1C), with the exception that participants were not asked to make a confidence judgment. Staircases for CFS and FBM target gratings were independent, and a threshold contrast value was estimated for each suppression method in each block (4 estimates total per suppression method). The median of these threshold contrast estimates for each suppression method was then multiplied by five different proportions, varied slightly from subject to subject by the experimenters, in order to target orientation discrimination performance values across the range of 60-90% correct, or, roughly speaking, d’ = 0.5-2.5. Multipliers used to determine FBM and CFS contrast values were as follows: Multipliers_FBM_ =0.57 ±0.08, 0.76 ±0.09, 0.94 ±0.10, 1.01 ±0.31, 1.30 ±0.13; Multipliers_CFS_ = 0.38 ± 0.09, 0.58 ± 0.07, 0.78 ± 0.11, 0.98 ±0.17, 1.19 ± 0.24. Notably, the multipliers used for CFS stimuli were lower than those used for FBM stimuli to account for the fact, which was gradually revealed to the experimenters as more participants were included, that the staircasing procedure had a greater tendency to overestimate threshold contrast values for CFS stimuli compared to FBM stimuli. Furthermore, to minimize potential ceiling effects that could arise from perceptual learning during the main experiment, staircasing threshold estimates over 75% contrast were excluded from the median threshold contrast calculation.

Additionally, if participants did not have threshold contrast estimates less than or equal to 75% contrast in at least two blocks for each suppression method, they repeated the same staircasing procedure (i.e., they performed an additional four staircasing blocks). If a participant repeated the staircasing procedure, threshold estimates from only the second staircasing procedure were used to determine the contrast values used in the main experiment, and threshold estimates up to 100% were included in the median threshold calculation. As long as a participant in the second staircasing procedure had at least one threshold contrast estimate under 100% for each suppression method, they were allowed to proceed to the main experiment. Otherwise, they were told that the experiment was finished and were excluded from participating in the main experiment. Five participants were excluded in this way. Notably, all five failed the QUEST procedure only for CFS stimuli, suggesting that the CFS task was, on average, more difficult to learn than the FBM task.

For the main experiment, a full factorial design was used in which all combinations of suppression method order (2), target orientation (2×2), and target contrast level for each suppression method (5×5) were presented, leading to a total of 200 unique trials. Each unique trial was presented twice, making for a total of 400 trials, which were randomized over eight 50-trial blocks. At the end of each block, participants were allowed to take a break with no time limit. At this time they were also given a score corresponding to their performance on the previous block, which was computed according to the following rules: one point was added or subtracted for each correct or incorrect orientation judgment, respectively. An additional point was either added or subtracted for each trial in which they correctly or incorrectly, respectively, discriminated the target orientation in the interval in which they indicated higher confidence. Participants were given a bonus of $10 USD if their final score exceeded that of the previous participant.

After participants completed the main experiment they were asked verbally by the experimenter whether, across the main experiment, they noticed any differences between the two stimulus intervals beyond basic differences in difficulty. This question was important in determining whether there may have been decisional or other cognitive response biases influencing subjects’ confidence responses. For example, if a participant could consistently distinguish between the FBM and CFS intervals, they might have consciously associated one of the two with higher confidence and, consequently, bet on that interval more frequently.

#### Data Analysis

The main question that was investigated in each of the current studies was whether or not we could find a difference in the relationship between subjective awareness and objective performance between two visual suppression methods. To get at this question, we used orientation discrimination d’ (Green & Swets, 1966) as an index of objective performance and confidence judgments as an index of subjective awareness (Lau & Rosenthal, 2011; Fleming & Lau, 2014).

For each subject, data were collapsed across target orientation order (Left-Left, Left-Right, Right-Left, Right-Right) and mask order (FBM-CFS, CFS-FBM) for each combination of contrast levels (5 FBM contrasts x 5 CFS contrasts = 25 combinations) in each trial. Orientation discrimination d’ was calculated for each suppression method for each of these contrast combinations. Type 1 hits were defined as trials in which the target had a left tilt and the subject chose left. Type 1 false alarms were defined as trials in which the target had a right tilt and the subject chose left. To adjust for values of infinite d’ in all experiments we used a standard correction that converts hit rates and false alarm rates of 1 and 0 to 1 - 1/2N and 1/2N, respectively, where N is the number of trials used in the calculation of d’ (MacMillan & Creelman, 2005).

We then plotted, for each of the 25 contrast combinations for each subject, the proportion of trials in which the CFS interval was rated with higher confidence against the difference in d’ between the CFS and FBM intervals (see Figure 2B). Individual psychometric curves were then generated by fitting the resulting 25 data points with a cumulative normal distribution function with free parameters α (threshold) and β (slope), and fixed parameters γ (lapse rate) = 0 and δ (guess rate) = 0, using the Palamedes Toolbox (Kingdom & Prins, 2009).

**Figure 2.**
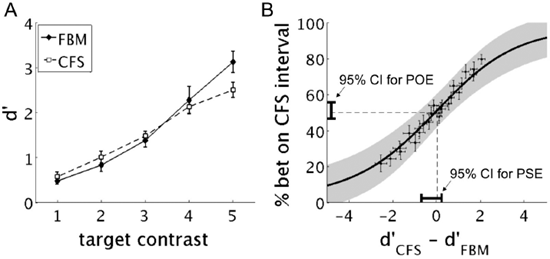
Results from Experiment **1. A)** Orientation discrimination performance (d’) at increasing target contrast under FBM (solid line) and CFS (dashed line). Error bars indicate ± **1** SEM. **B)** Average psychometric curve. For each participant, the proportion of trials in which they bet on the CFS interval was plotted as a function of the difference in d’ between CFS and FBM intervals for each of the 25 combinations of stimulus contrast levels that could occur in a single trial (shown are group means ± **1** SEM). A cumulative normal function was then fit to each participant’s data, with mean and slope as free parameters. Plotted is the mean of the individual participant fits (black line) ± **1** SD (gray). The 95% confidence interval for the estimated PSE and POE group means are shown by the black bars sitting near the x- and y-axes, respectively. A significant rightward shift of the psychometric curve, such that the confidence interval for the PSE were to fall above of zero, would suggest that when subjective awareness is matched between CFS and FBM, d’ is significantly higher under CFS than under FBM. Similarly, if a rightward shift of the psychometric curve makes it such that the confidence interval for the POE falls below 50%, then it would suggest that subjective awareness of the target stimulus is higher under FBM when d’ is matched. This would indicate relative blindsight. The opposite interpretations would hold if the confidence interval for PSE were below zero and the confidence interval for POE were above 50%. The fact that zero falls within the observed PSE confidence interval suggests that when subjective awareness of the target was matched between CFS and FBM, there was no significant difference in discrimination d’ between the two suppression methods. Similarly, the fact that 50% falls within the observed POE confidence interval suggests no evidence for relative blindsight.

If there is no difference in the relationship between subjective awareness and objective performance between two given suppression methods, then we should expect the point of subjective equality (PSE), or the difference in d’ at which participants are equally likely to bet on the two suppression methods, to be zero. Similarly, the point of objective equality (POE), or the likelihood of betting on the CFS interval when d’_CFS_ - d'_FBM_ = 0, should be 50%. If, on the other hand, the relationship between subjective awareness and objective performance is significantly different between the two suppression methods, then the psychometric function should shift such that the PSE and POE should be significantly different from zero and 50%, respectively. Therefore, in each of the following experiments, the main tests of interest were one-sample t-tests (α = .05, two-tailed) conducted on the PSEs and POEs obtained from the individually-fitted psychometric functions, with the null hypothesis being that the mean PSE and POE across participants are equal to zero and 50%, respectively. In the case of the POE analysis, since d’ is matched between suppression methods, subjective awareness is operationally defined in this case in line with Giles, Lau, & Odegaard (2016), as the difference in Type 2 responding when Type 1 performance is matched. Analyses for each experiment were conducted in Matlab R2013a (Natuck, MA), with the exception of repeated measures ANOVAs, which were conducted in SPSS v22 (IBM, Armonk, NY, USA). All repeated measures ANOVAs were adjusted for violations of the assumption of sphericity with the Greenhouse-Geisser correction when necessary (Experiments 1 and 2 only).

### Results & Discussion

A repeated measures ANOVA with within-subjects factors contrast (5 levels) and suppression method (FBM or CFS) revealed the expected main effect of contrast on orientation discrimination d’ [F(2.29,41.25) = 75.97, p < 0.001] (Figure 2A), i.e. that increased contrast led to higher performance. The ANOVA also showed no main effect of suppression method [F(1,18) = 0.52, p = 0.48], but a significant interaction between contrast and suppression method [F(2.33,42.00) = 4.59, p = 0.012]. Figure 2A suggests that this interaction is driven by the sudden divergence in d’ between suppression methods at the highest contrast level. This was confirmed by post hoc Bonferroni corrected two-tailed paired t-tests [at contrast level 5: t(18) = 3.82, p = 0.001, whereas p-values for contrast levels 1-4 were all > 0.38]. Because, by design, contrast levels for the main experiment were selected on a subject-by-subject basis with the goal of optimally matching d’ between suppression methods, this result is mostly attributable to experimenter error. Nonetheless, the lack of a main effect of suppression method on discrimination d’ in the ANOVA indicates that, across contrast levels, Type 1 performance was matched between FBM and CFS. However, to check for any potential biasing of the PSE and POE analyses that could have resulted from the difference in d’ between suppression methods at the highest contrast level, we conducted the PSE and POE analyses once with all contrast levels included, and once using only contrast levels 1-4.

As for the main analyses, looking across all contrast levels, PSE and POE values were −0.25 ± 0.22 and 51% ± 2%, respectively. Two-tailed paired t-tests indicated insufficient evidence to reject the null hypotheses that the PSE is equal to zero [t(18) = −1.14, p = 0.27, 95% CI = (−0.70, 0.21)] and the POE is equal to 50% [t(18) = 0.53, p = 0.60, 95% CI = (47%, 56%)] (Figure 2B). When we excluded the highest contrast level, there was still insufficient evidence to reject the null hypothesis in each case [PSE: t(18) = −0.95, p = 0.35, 95% CI = (−0.71, 0.26); POE: t(18) = 0.82, p =0.42, 95% CI = (47%, 56%)]. The PSE results suggest that when subjective awareness is matched, there is no difference in the level of objective performance under FBM and CFS. Similarly, the POE analysis suggests that when Type 1 performance is matched, there is no difference in participants’ subjective awareness of the target stimulus between the two suppression methods.

Importantly, all participants responded in the negative when asked, after the main experiment, if on any trials they noticed differences between the two intervals beyond difficulty level. This suggests that participants’ confidence judgements were not subject to decisional biases based on explicit knowledge about the difference between FBM and CFS stimuli.

One concern is that the gaps between masks in the CFS condition, which lead to a collective 156.9 ms in which the target is presented to one eye with no mask presented to the other eye (Figure 1B), may minimize the degree to which the CFS condition elicits a true binocular rivalry effect. If this is the case, then, presumably, it should also minimize mechanistic differences underlying the disruption of visual processing between the two suppression methods, thereby reducing our chances of rejecting the null hypothesis.

One potential piece of evidence that FBM and CFS use different mechanisms to disrupt visual processing is that target contrast values were significantly lower for CFS stimuli (24.15 ± 3.03% Michelson contrast) than they were for FBM stimuli (36.89 ± 1.14% Michelson contrast) [two-tailed, paired t-test: t(18) = 4.66, p < 0.001]. Importantly, this result holds when excluding the highest contrast level [two-tailed, paired t-test: t(18) = 4.87, p < 0.001]. However, an alternative interpretation is that the lower contrast thresholds found in the CFS condition are simply driven by the longer presentation times for CFS target stimuli relative to FBM target stimuli. Disambiguating these hypotheses is critical for establishing that the FBM and CFS conditions induce mechanistically different visual suppression effects. We address this issue directly in Experiment 2.

## Experiment 2

### Methods

#### Participants

Nine participants (3 female, ages 18-39, 3 left-handed, 4 left-eye dominant), including the first author, gave written informed consent to participate. Seven of the nine participants in Experiment 2 had also participated in Experiment 1. One participant (inexperienced) was excluded due to reporting incomplete fusion of binocular stimuli on many trials during the main experiment. Therefore, 8 total participants (2 female, ages 21-39, 2 left-handed, 4 left-eye dominant, 1 inexperienced) were included in the analyses for Experiment 2. All participants had normal or corrected-to-normal eyesight and normal stereo vision, and all were either paid $10 USD or given course credit for their participation.

#### Apparatus and Stimuli

Apparatus and stimuli in Experiment 2 were the same as in Experiment 1, except for the following. Instead of CFS, we used a binocular rivalry technique conventionally referred to as interocular suppression (IS) (Breitmeyer, 2015; Izatt et al., 2014). The sole difference between the CFS condition in Experiment 1 and the IS condition in Experiment 2 is that the target grating no longer had its contrast ramped up from and down to zero. Instead, the IS target grating had the same duration as the FBM target grating (26.7 ms), and its contrast was constant (Figure 3A). Furthermore, in each interval the target had an equal probability of appearing between either masks 2 and 3 or masks 3 and 4. The randomization was independent in the two intervals such that in approximately half of all trials (48% ± 3%) the target appeared between the same mask numbers (e.g., 2 and 3) in each interval, while in the remainder of trials the target appeared between different mask numbers in each interval (e.g., between masks 2 and 3 for FBM and masks 3 and 4 for IS). This manipulation was introduced to minimize the degree to which participants could anticipate the timing of target onset. Such anticipation, whether conscious or unconscious, could potentially minimize visual processing differences between the two masking conditions. Three participants shifted the left fusion contours at the beginning of the experiment by 2.39° ± 1.63° leftward and 0.01° ± 0.25° upward.

**Figure 3.**
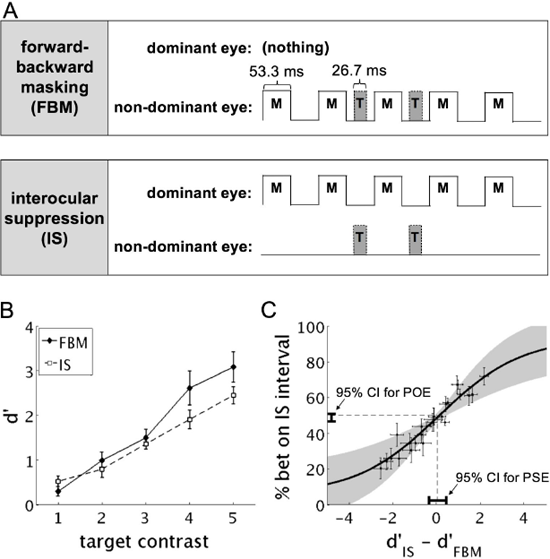
Stimuli and results from Experiment 2. **A)** Temporal dynamics of stimuli from Experiment 2. Mask stimuli had the same temporal profile as those in Experiment 1. Target stimuli in the IS interval appeared abruptly at peak contrast instead of ramping up and down in contrast as in Experiment 1. In each interval target stimuli were presented pseudo randomly between either the second and third or third and fourth masks. **B)** Orientation discrimination performance (d’) at increasing target contrast under FBM (solid line) and IS (dashed line). **C)** Average psychometric curve and 95% confidence intervals for estimated PSE and POE group means, calculated and shown the same way as in Experiment 1 (see methods, Figure 2). Because the PSE confidence interval contains the point d’_difference_ = 0 and the POE confidence interval contains the point at which subjects were 50% likely to bet on either suppression method, these results suggest that there was no evidence for a difference in the relationship between objective and subjective thresholds between FBM and IS. Error bars in B & C indicate ± 1 SEM. Gray region in C indicates ± 1 SD of psychometric fits.

Therefore, across the entire experiment, the only difference between FBM and IS stimuli was ocularity (Figure 3A). It follows that if we observe differences in contrast thresholds and stimulus contrast values at matched d’ between FBM and IS similar to those found between FBM and CFS in Experiment 1, then these differences should be attributed to the difference in ocularity between the two conditions. This result would provide additional evidence for the presence of a binocular rivalry-based suppression effect in our original CFS condition.

We also reasoned, based on previous evidence for a higher degree of subliminal priming under FBM than IS (Izatt et al, 2014; Breitmeyer, 2015), that IS may allow a greater degree of unconscious orientation discrimination than FBM. If true, we would expect a leftward shift in the psychometric function such that at the PSE there would be significantly higher discrimination d’ under FBM than under IS, and at the POE there would be a significantly higher tendency to bet on the IS interval.

#### Procedure and Data Analyses

The procedure in Experiment 2 was the same as that in Experiment 1 except for the following. Different proportions of the median threshold contrast estimate from staircasing were used to determine target contrast values for the main experiment (Multipliers_FBM_ = 0.58 ±0.12, 0.75 ± 0.13, 0.93 ±0.14, 1.11 ±0.16, 1.29 ±0.17; Multipliers_IS_ = 0.36 ± 0.07, 0.57 ± 0.04, 0.79 ± 0.09, 1.01 ± 0.15, 1.23 ± 0.21). Again, the multipliers used for IS were lower than those used for FBM to account for the tendency of the staircasing procedure to overestimate threshold contrast values to a greater extent for IS stimuli than for FBM stimuli.

Additionally, 40 catch trials, in which the contrast of the target grating in one of the two intervals (counterbalanced between suppression methods) was at 100%, were randomly interleaved among the 400 main experiment trials. This made for a total of 440 trials in the main experiment, which were divided into eight 55-trial blocks. These catch trials were added both to help participants maintain perceptual templates of the left- and right-tilted target gratings, and to keep participants motivated throughout what is otherwise a very difficult and, according to anecdotal evidence from some participants following Experiment 1, sometimes demoralizing task.

Analysis procedures followed those conducted in Experiment 1.

### Results & Discussion

A repeated measures ANOVA with within-subjects factors contrast (5 levels), suppression method (FBM or IS), and target timing (between masks 2 and 3 or between masks 3 and 4) again showed the expected main effect of contrast on orientation discrimination d’ [F(1.72,12.07) = 42.57, p < 0.001; Figure 3A]. As in Experiment 1, there was no main effect of suppression method [F(1,7) = 2.22, p = 0.18]. Unlike Experiment 1, however, there was no interaction between contrast and suppression method [F(1.90,13.26) = 0.94, p = 0.41], suggesting that discrimination d’ was matched effectively between the two suppression methods across contrast levels.

Interestingly, there was a main effect of target timing on discrimination d’ [F(1,7) = 15.55, p = 0.006]. A post hoc two-tailed paired t-test on discrimination d’ values calculated across suppression methods and contrast levels showed that d’ was significantly higher when the target stimulus was presented between masks 2 and 3 than when it was presented between masks 3 and 4 [t(7) = 3.73, p = 0.007]. There was no interaction between target timing and contrast [F(2.51,17.60) = 0.44, p = 0.69], however, there was a significant interaction between target timing and suppression method [F(1,7) = 6.62, p = 0.037]. Post hoc Bonferroni-corrected two-tailed paired t-tests showed that d’ was significantly higher when the target was presented earlier under FBM [t(7) = 3.65, p = 0.008], but that there was only a marginal trend towards this relationship under IS [t(7) = 2.02, p = 0.08]. There was no significant 3-way interaction [F(2.83,19.83) = 0.44, p = 0.71].

The effect of stimulus timing on objective performance may be attributable to rhythmic attentional sampling (Landau & Fries, 2012) set by visual cues preceding the onset of the target stimulus (e.g., the initial fixation cross or the onset of the first mask). The difference in this effect between FBM and IS conditions may provide evidence for a functionally relevant difference in the mechanism of visual suppression between the two suppression methods.

As in Experiment 1, we did not find evidence to reject the null hypothesis that d’ is matched between suppression methods at the PSE [t(7) = 0.23, p = 0.82, 95% CI = (−0.36, 0.44); Figure 3B], Similarly, we did not find evidence to reject the null hypothesis that subjects are equally likely to bet on each suppression method at the POE (i.e., there was no evidence for relative blindsight) [t(7) = −1.44, p = 0.19, 95% CI = (46%, 51%)]. It was, however, verified that target contrast values were again lower under IS (14.96 ± 2.51%) than they were under FBM (35.94 ± 3.12%) [t(7) = 9.82, p < 0.001]. This provides additional evidence for a difference in the mechanism of visual suppression between FBM and the binocular conditions in both Experiments 1 and 2, despite the absence of the hypothesized difference in the relationship between objective performance and subjective awareness.

Also consistent with Experiment 1, no participants indicated noticing a difference between FBM and IS intervals when questioned after the main experiment. Furthermore, participants were 100% correct when discriminating catch trial target stimuli with 100% contrast. Betting accuracy on catch trials was similarly high (98.75 ± 0.82% correct, where a correct bet is defined as a bet on an interval in which the orientation judgment was correct), suggesting that participants were maintaining attention throughout the experiment.

Given the failure to reject the null hypothesis in the first two experiments, we next turned to backward masking (BM) as an alternative to FBM. Previous evidence suggests that BM, but not CFS, allows for the subliminal priming with non-manipulable objects (Almeida et al., 2008). It has also been suggested that, relative to FBM, the visual signal under BM may benefit from an increased signal-to-noise ratio when performance is matched (Breitmeyer, 2015; Harris et al., 2011; Macknik & Livingstone, 1998). We therefore hypothesized that BM may allow for a greater degree of unconscious processing than CFS, and that, in our 2IFC paradigm, we may therefore see the psychometric function shift so as to show higher discrimination d’ under BM at the PSE, and a higher tendency to bet on the CFS interval at the POE.

## Experiment 3

### Methods

#### Participants

Eight participants (3 female, ages 22-39, 2 left-handed, 5 left-eye dominant), including the first author, gave written informed consent to participate. Six of the eight participants in Experiment 3 had also participated in Experiment 1, and five of these participants had also participated in Experiment 2. All participants had normal or corrected-to-normal eyesight and normal stereo vision, and all were either paid $10 USD or given course credit for their participation.

#### Apparatus and Stimuli

Apparatus and stimuli in Experiment 3 were the same as in Experiment 1, except for the following. For both BM and CFS conditions, mask stimuli were shifted later in time by 26.7 ms.

In the BM interval, the first mask was preceded by the target, which had a duration of 26.7 ms, meaning target offset coincided with mask onset. In the CFS interval, target onset coincided with the onset of the first mask and returned to the same ramping dynamics used in Experiment 1 (Figure 4A). Only one participant shifted the left fusion contours at the beginning of the experiment (3.44° leftward with no vertical shift).

**Figure 4.**
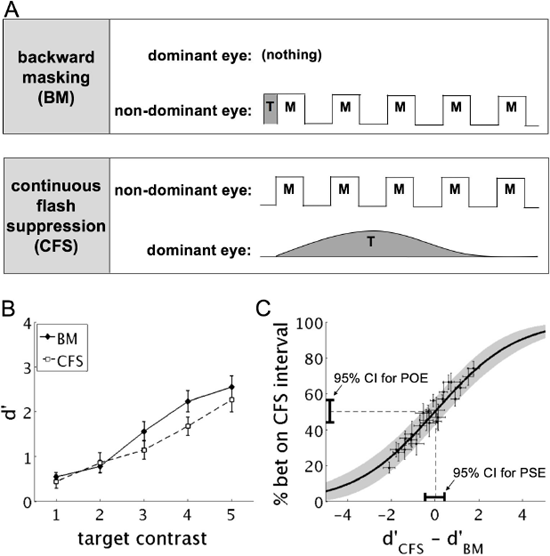
Stimuli and results from Experiment 3. **A)** Temporal dynamics of stimuli from Experiment 3. In BM intervals the target was first presented for 26.7 ms and was immediately followed by the first mask. Five masks were presented for 53.3 ms each, with intervening blank gaps of the same length. The offset of the last mask was followed by a blank gap of 26.7 ms. Masks in the CFS interval had the same temporal profile as those in the BM interval. The onset of the target stimulus in the CFS interval occurred simultaneously with the onset of the first mask and otherwise had the same temporal ramping profile as the target stimulus in Experiment 1 (see methods, Figure 1). **B)** Orientation discrimination performance (d’) at increasing target contrast under BM (solid line) and CFS (dashed line). **C)** Average psychometric curve and 95% confidence intervals for estimated PSE and POE group means, calculated and shown the same way as in Experiment 1 (see methods, Figure 2). Because the PSE confidence interval contains the point d’_difference_ = 0 and the POE confidence interval contains the point at which subjects were 50% likely to bet on either suppression method, these results suggest that there was no evidence for a difference in the relationship between objective and subjective thresholds between BM and CFS. Error bars in B & C indicate ± 1 SEM. Gray region in C indicates ± 1 SD of psychometric fits.

#### Procedure

The procedure in Experiment 3 was the same as that in Experiment 2 except for the use of different proportions of the median threshold contrast estimate to determine target contrast values for the main experiment (Multipliers_BM_ = 0.23 ± 0.07, 0.40 ± 0.06, 0.57 ± 0.05, 0.77 ± 0.07, 0.99 ±0.12; Multipliers_CFS_ = 0.32 ± 0.11, 0.50 ± 0.08, 0.68 ± 0.06, 0.87 ± 0.05, 1.10 ± 0.08). Interestingly, median threshold contrast estimates were, on average, more overestimated under BM than they were under CFS. As a result, the multipliers used to determine target contrast levels for the main experiment were lower for BM stimuli than they were for CFS stimuli.

### Results & Discussion

Consistent with Experiments 1 and 2, a repeated measures ANOVA with within-subject factors contrast (5 levels) and suppression method (BM or CFS) showed the expected main effect of contrast [F(4,28) = 96.20, p < 0.001; Figure 4B] and no main effect of suppression method [F(1,7) =1.09, p 0.33]. Consistent with Experiment 2, there was no significant interaction between contrast and suppression method [F(4,28) = 1.20, p = 0.33], again suggesting that d’ was effectively matched between suppression methods across contrast levels.

Regarding the main analysis, once again there was not sufficient evidence to reject the null hypothesis that d’ is matched between BM and CFS at the PSE [t(7) = 0.20, p = 0.85, 95% CI = (−0.47, 0.40); Figure 4C]. Nor was their sufficient evidence to reject the null hypothesis that subjects are equally likely to bet on each suppression method at the POE [t(7) = 0.12, p = 0.91, 95% CI = (44%, 57%)], again providing no evidence for relative blindsight. Experiment 3 was therefore in line with Experiments 1 and 2 in providing no evidence for a difference in the relationship between objective performance and subjective awareness between suppression methods.

Interestingly, in contrast to Experiments 1 and 2, mean stimulus contrast per subject in the main experiment was not significantly different between BM (11.39 ± 2.85%) and CFS (18.87 ± 4.42%) [t(7) = 1.87, p = 0.10]. This decrease in threshold target contrast from FBM to BM is presumably due to the relative lack of interference with feedforward processing under BM (Breitmeyer, 2015; Harris et al.,2011; Macknik & Livingstone, 1998).

No participants reported noticing a difference between BM and CFS intervals when questioned after the main experiment. Performance on catch trials was again high (orientation judgment accuracy: 97.81 ± 1.86% correct, betting accuracy: 98.13 ± 1.03% correct), suggesting that participants were maintaining attention throughout the task.

## General Discussion

In three experiments we looked for a difference in the relationship between objective performance and subjective awareness, in line with reports of relative blindsight (Lau & Passingham, 2006), between pairs of visual suppression methods. In each case we found no evidence for any such difference, suggesting that the relationship between objective and subjective thresholds for forced-choice orientation discrimination is equivalent under FBM, CFS, IS, and BM. Taking a specific definition of subjective awareness (Giles et al., 2016), which is operationally defined as what is tracked by subjective reports while sensitivity is controlled for, we interpret the results (i.e. the POE analyses) to mean that the different suppression methods impact subjective awareness similarly.

We used a modified version of the 2IFC paradigm from Peters & Lau (2015) in which each of two suppression methods, one per 2IFC interval, was used to mask a left- or right-tilted target grating. Subjective awareness was indexed by forcing participants to bet on the interval in which they had higher confidence in their ability to discriminate the orientation of the target grating. This paradigm has several advantages that build on previous studies comparing different visual suppression techniques. For instance, some studies have compared suppression techniques *between* experiments (Almeida et al., 2008, 2010, 2013; Faivre, Berthet, & Kouider, 2012), making them vulnerable to potentially confounding idiosyncratic differences between experimental conditions. Further, the forced-choice nature of the subjective judgment reduces concern about subjective criterion biases that may have been present in previous comparative suppression studies (Izatt et al., 2014; Peremen & Lamy, 2014). To further reduce subjective biases, we took inspiration from earlier studies that compared monocular and binocular suppression conditions within single experiments (Jiang et al., 2007; Stein et al., 2011; Izatt et al., 2014) and designed stimuli such that, beyond simple differences in difficulty, the two intervals on a given trial appeared subjectively similar. This has the benefit of minimizing conscious decisional biases (e.g., participants having a conscious preference for backward masked stimuli over CFS-masked stimuli) that would otherwise reduce the chances of finding the hypothesized difference in the relative positioning objective and subjective discrimination thresholds between suppression methods.

We interpret these findings to suggest, in line with Peters & Lau (2015), that objective and subjective thresholds under these conditions do not dissociate. That is to say, we consider the current results to be further evidence against the idea that normal observers have any capacity for unconscious orientation discrimination. This idea is in line with others who have argued that objective thresholds should, a priori, be considered equivalent to subjective thresholds in forced-choice perceptual tasks (Snodgrass & Shevrin, 2006; Phillips, 2017). These findings further suggest that controlling for criterion bias may be a critical experimental difference between studies that report evidence for unconscious forced-choice discrimination sensitivity (Lamy, Salti, & Bar-Haim, 2008; Hesselman et al., 2011; Salti et al., 2015) and those that report evidence against it (Peters & Lau, 2015).

An important limitation is that it remains an open question whether a different visual suppression technique can selectively impair subjective awareness while leaving objective discrimination performance relatively intact. Future studies should compare visual suppression techniques that are more distant from each other in terms of how much unconscious priming they allow, e.g., FBM and visual crowding (Breitmeyer, 2015), or that have been functionally characterized to act at different points in the visual processing stream, e.g., visual crowding and object substitution (Chakravarthi & Cavanagh, 2009) or metacontrast masking and interocular suppression (Breitmeyer et al., 2008). They can also focus on suppression methods that rely on attentional manipulations (e.g., attentional blink, inattentional blindness), which may allow for higher levels of unconscious processing (Kouider & Dehaene, 2007) that include unconscious forced-choice discrimination. The current paradigm provides a useful means for comparing such suppression techniques, while maintaining a rigorous control for criterion bias. However, a challenge in designing these studies will be in creating stimuli that make the techniques under comparison appear superficially indistinguishable.

It should also be emphasized that we extend the current interpretation of a lack of unconscious perception only to *direct* perceptual tasks such as forced-choice detection and discrimination tasks (Green & Swets, 1966; Macmillan & Creelman, 2004), and not to other established indirect perceptual effects like subliminal priming (Hannula, Simons, & Cohen, 2005; Kouider & Dehaene, 2007; though see Phillips (2017) for a discussion on whether priming effects should constitute genuine cases of perception per se). Even if we assume that normal observers do have some capacity for direct unconscious perception, our results suggest that we should not expect hierarchical relationships for subliminal priming among suppression methods (e.g., Kouider & Dehaene, 2007; Faivre, Berthet, & Kouider, 2014; Breitmeyer, 2015) to apply to direct unconscious perception. For example, Almeida et al. found greater subliminal priming effects for tool stimuli (2008, 2010) and emotional faces (2011) under BM than under CFS, while Izatt et al. (2014) found greater subliminal face priming effects under FBM than under IS. These hierarchical relationships among suppression methods for subliminal priming clearly conflict with the null results for differences in direct unconscious processing between suppression methods observed here. However, even some previously suggested hierarchical relationships between suppression methods should be approached with caution, as judgments of prime visibility in these studies were vulnerable to criterion bias (Izatt et al., 2014; Peremen & Lamy, 2014). The 2IFC paradigm described in Peters & Lau (2015) provides a means for future priming studies to ensure invisibility of primes without this potential confound.

Another limitation in both the current study and Peters & Lau (2015) is that participants may have ignored instructions to rate confidence specifically in their performance on the orientation discrimination task, and instead rated confidence based on the *detectability* of target stimuli. In both studies, exclusive use of such a heuristic would lead to the observed null results. A major difference between the two studies is that in the current study, a valid target grating was presented in both intervals on every trial, whereas in Peters & Lau (2015), each trial contained exactly one interval in which the target grating was completely absent. We might expect, therefore, that participants in the current study would have been less motivated to use a detection heuristic, as the higher proportion of trials in which detectability is matched should make target detectability less automatically informative for rating confidence. It should be noted, however, that the current design does not eliminate the possibility that participants used such a detection heuristic anyway. Indeed, previous evidence from studies using orientation discrimination tasks suggests that we should expect this to be the case (Koizumi, Maniscalco, & Lau, 2015; Maniscalco, Peters, & Lau, 2016). Future studies could test this question directly, for example, by using a 2IFC design in which one interval contains a valid target and the other contains an uninformative, but stimulus energy-matched target (e.g., a vertical grating).

Importantly, an alternative interpretation of our main result is that all of the suppression methods used in the current study allowed an equal, greater-than-zero amount of direct unconscious perception. We argue against this interpretation based on the findings of Peters & Lau (2015), which provide evidence for a lack of direct unconscious perception under FBM. Based on this, and given that no difference in the relationship between objective and subjective thresholds was observed between FBM and any of the other suppression techniques used here (whether tested directly or implied by transitive logic), we conclude that no direct unconscious perception occurred under any of the present suppression techniques. If objective and subjective thresholds do actually dissociate under FBM (contra the conclusion of Peters & Lau (2015)) –- for example, because subjects use a detection heuristic that prevents unconscious perception from being detected via the 2IFC method –- then the current data suggest that the three other suppression techniques used here cause this dissociation *to precisely the same extent.* While this is theoretically tenable, we argue that it seems less parsimonious than the alternative interpretation that each suppression method simply fails to cause a dissociation between objective and subjective thresholds at all. As mentioned, this interpretation is in line with previous arguments that any direct perceptual discrimination sensitivity should coincide with some degree of perceptual consciousness (Snodgrass & Shevrin, 2006; Phillips, 2017; but see Block in Phillips & Block, 2016 for an opposing view).

In conclusion, we have shown a lack of a difference in the relationship between objective and subjective thresholds for forced-choice orientation discrimination between four commonly used visual suppression techniques. Taken together with previous evidence (Peters & Lau, 2015), these results suggest that when criterion bias is sufficiently controlled for, normal observers do not demonstrate direct unconscious perception. Whether this capacity can be demonstrated under a different set of visual suppression conditions is a matter for future studies to investigate. The present results should, however, place helpful constraints on future hypotheses and methodological choices for studying conscious and unconsciousness visual perception.

## Acknowledgements

**Acknowledgments.** This work was supported by a grant from the National Institutes of Health (US) to HL (grant number R01NS088628), and a National Science Foundation Graduate Research Fellowship to JDK.

## Data and program code availability

Data and program code from this study are available from the corresponding author upon reasonable request.

